# MetaXtract: Extracting Metadata from Raw Files for FAIR Data Practices and Workflow Optimisation

**DOI:** 10.1101/2025.11.12.687968

**Authors:** Ahmad Lutfi, Zhuo A. Chen, Lutz Fischer, Juri Rappsilber

## Abstract

Mass spectrometry (MS) experiments generate rich acquisition metadata that are essential for reproducibility, data sharing, and quality control (QC). Because these metadata are typically stored only in vendor-specific formats, they often remain difficult to access. MetaXtract is a lightweight tool that extracts detailed parameters directly from Thermo Fisher raw files and exposes them in structured, tabular formats. By capturing sample information, LC-MS method settings, and scan-level metrics such as retention time, total ion current, and ion injection time, MetaXtract increases transparency and ensures that essential acquisition details accompany published data and results in easy readable form. This supports FAIR data practices by improving the findability, accessibility, interoperability, and reusability of MS datasets after converting them to other formats, thereby increasing the value of deposition in public repositories. The importance of such metadata accessibility was recently highlighted by the crosslinking mass spectrometry community in efforts to advance FAIR data principles, and it extends to MS-based omics approaches more broadly. Importantly, MetaXtract enables search-free, near real-time performance monitoring by relying on acquisition-side signals, providing actionable indicators immediately after data acquisition rather than after database searching. This also caters for laboratory or depository internal streamlined QC and troubleshooting through integration into automated pipelines. By embedding acquisition parameters into routine data handling, MetaXtract strengthens reproducibility, optimises method development, and supports large-scale applications, including machine learning and secondary data analysis.

**Graphical Abstract:** 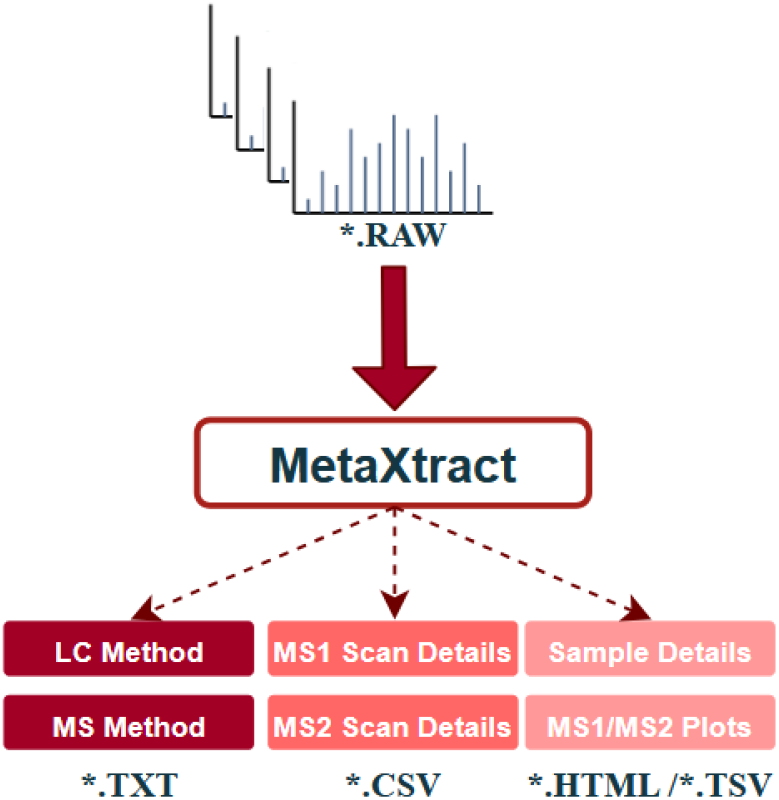

**Highlights:**

- Metadata extraction from Thermo Fisher *raw* files
- Enhanced findability, accessibility, interoperability, and reusability of deposited data
- Integration into workflows via GUI and command-line modes
- Troubleshooting support by visualizing MS1/MS2 scan details
- Indexed MS1/MS2 peak list export enabling machine learning workflows

**Availability:** MetaXtract is available for free download as open-source software at https://github.com/Rappsilber-Laboratory/MetaXtract, the software is licensed under the Apache-2.0 license.

## Introduction

Mass spectrometry (MS) generates large binary raw files containing both spectral data (MS1 and MS2) and essential acquisition metadata such as precursor charge states, retention times, ion injection times, and fragmentation energies (1). Although these parameters are critical for data interpretation, reproducibility, and quality control, they are cumbersome to access at scale because vendor-specific formats typically require proprietary software. Beyond supporting reproducibility and reporting, acquisition metadata provide instrument- and method-level signals that can diagnose technical variance independently of peptide/protein identifications. In Thermo RAW files, relevant parameters are distributed across scan headers, instrument methods, and auxiliary records, and are typically exposed only through vendor APIs that can be platform-limited and inconsistent across software/instrument generations. This makes standardized, batch-scale extraction and downstream integration difficult in routine workflows. Existing vendor tools offer only limited flexibility for automated extraction, batch processing, or integration into modern bioinformatics pipelines. Consequently, researchers lack practical means to obtain comprehensive and structured metadata for QC, method optimisation, and large-scale computational workflows (2, 3), highlighting the need for more accessible and robust extraction solutions (1).

Recent community efforts to standardise crosslinking MS have highlighted the importance of complete and machine-readable metadata to achieve FAIR data practices and reproducibility. The Crosslinking MS Roadmap (4) emphasises that, while mzIdentML 1.3 provides a unified format for reporting identified crosslinks (5, 6), critical acquisition parameters such as LC-MS method settings and scan-level metrics remain largely inaccessible. Extracting these parameters directly from mass-spectrometric raw data is considered essential for automated repository submission and transparent provenance tracking. Addressing this community-defined need across proteomics demands practical tools capable of reliably capturing such metadata and providing it in structured, reusable formats.

Although several tools have been reported to offer access to some metadata, these existing tools present significant limitations (**Table 1**). RawMeat (7), developed by Vast Scientific, is no longer supported, making it impractical for long-term use. The MSQC software (8, 9), developed by the National Institute of Standards and Technology (NIST), facilitates QC by computing metrics from LC-MS/MS data, but it requires database search results for most metrics, limiting real-time instrument monitoring. RawBeans (10) lacks flexibility in selecting MS1/MS2 data and does not extract LC-MS method information, complicating method validation and troubleshooting. Its lack of updates also raises compatibility concerns with newer instruments. LogViewer (11) depends on log-files from RawXtract (.*ms2* and .*log* files), making it unsuitable for high-throughput workflows. It also does not extract method settings and is no longer available for download. SIMPATIQCO (12) has not been updated since 2015, cannot export selected metadata or QC metrics, and requires advanced user expertise for setup and use. Similarly, QuaMeter (13) is no longer accessible from its official source and lacks ongoing support, limiting its use in current environments. In general, these limitations point to the need for a modern and robust QC solution in proteomics: outdated software, lack of flexibility, missing method information, and download/accessibility issues.

**Table 1.**
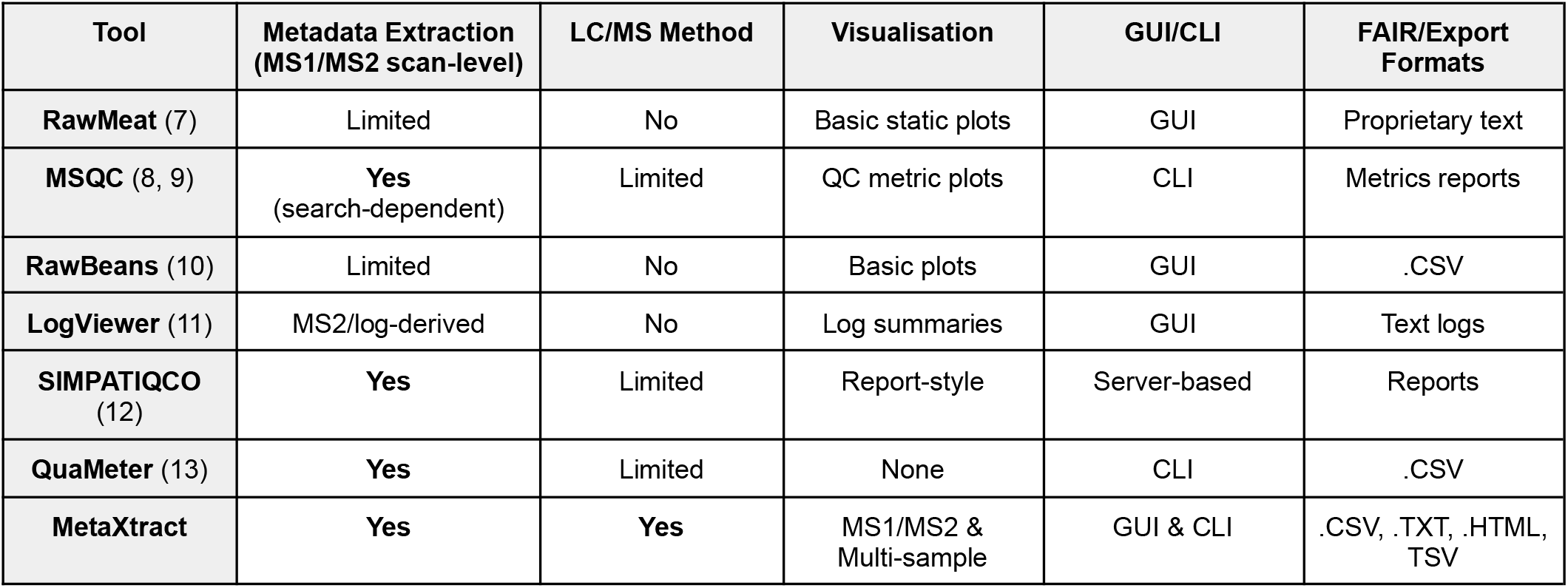
Comparative features relevant to QC, metadata extraction, and FAIR export in LC-MS/MS tools, including MetaXtract. *GUI: Graphical User Interface; CLI: Command Line Interface*.

| Tool | Metadata Extraction<br>(MS1/MS2 scan-level) | LC/MS Method | Visualisation | GUI/CLI | FAIR/Export<br>Formats |
| --- | --- | --- | --- | --- | --- |
| **RawMeat** (7) | Limited | No | Basic static plots | GUI | Proprietary text |
| **MSQC** (8, 9) | **Yes**<br>(search-dependent) | Limited | QC metric plots | CLI | Metrics reports |
| **RawBeans** (10) | Limited | No | Basic plots | GUI | .CSV |
| **LogViewer** (11) | MS2/log-derived | No | Log summaries | GUI | Text logs |
| **SIMPATIQCO**<br>(12) | **Yes** | Limited | Report-style | Server-based | Reports |
| **QuaMeter** (13) | **Yes** | Limited | None | CLI | .CSV |
| **MetaXtract** | **Yes** | **Yes** | MS1/MS2 &<br>Multi-sample | GUI & CLI | .CSV, .TXT, .HTML,<br>TSV |

The rapid expansion of mass spectrometry based proteomics over the past decade is reflected by the steep increase in RAW file submissions to PRIDE (Supplementary Figures S1-S2). With about 75 % of submitted MS files (for the years from 2000 to 2025 consult Supplementary Data), Thermo RAW files overwhelmingly dominate annual submissions, consistently representing the largest proportion of deposited primary MS data, while other vendor formats such as Bruker .d and Sciex WIFF/WIFF2 contribute comparatively smaller fractions. This clear format asymmetry justifies our initial focus on native Thermo RAW support, as it maximizes immediate impact and coverage across publicly available datasets. By prioritizing the predominant acquisition format, the tool addresses the largest segment of real-world proteomics data while establishing a scalable framework for future community-based extension to additional vendor formats.

A practical solution should (i) export metadata into structured, analysis-ready tables, (ii) support both interactive inspection and automated batch processing, and (iii) provide immediate visualization for performance monitoring and troubleshooting without requiring identification results. To overcome the current limitations of QC tools, we developed MetaXtract, a lightweight Python-based tool that directly reads RAW files using the Thermo Fisher RAWFileReader library (14). It offers comprehensive MS1 and MS2 visualizations, allowing users to explore key metrics like precursor intensity, ion injection time, charge states, and retention time distributions. MetaXtract provides customizable tabular output, enabling users to select specific parameters. It also extracts sample information and LC-MS method details, supporting quality control and reproducibility.

In addition to metadata extraction, MetaXtract exports MS2 spectra as indexed lists (m/z, intensity, resolution, noise, charge state) linked to their corresponding scan-level metadata via scan numbers. MS1 spectra are exported analogously (m/z and intensity arrays). The output is written in columnar Parquet format, enabling memory-efficient access without loading entire files into memory and facilitating scalable downstream processing. Because spectral peak arrays remain directly linked to acquisition parameters such as retention time, precursor m/z, charge, injection time, and fragmentation settings, the resulting data structure is well suited for data-driven modelling approaches. For example, the exported data can be used to train models for spectral quality assessment, retention time prediction, fragmentation behaviour analysis, anomaly detection for instrument monitoring, or representation learning on raw spectral profiles with the possibility to import the tool into Python scripts.

With a user-friendly GUI, researchers can easily configure the outputs without advanced computational expertise. The tool can be integrated into workflow-based frameworks like Snakemake (15) and Nextflow (16) for efficient metadata extraction. MetaXtract installer and source code are available for free at https://github.com/Rappsilber-Laboratory/MetaXtract.

## Results and discussion

### MetaXtract Interface and Visualization Features

MetaXtract provides a GUI and a CLI. The tool can be downloaded from GutHub and runs using the source code or the installer (PyInstaller), simplifying its use also in managed desktop or otherwise restricted work environments. The GUI offers an intuitive interface for browsing files, inspecting scan measurements, and generating plots interactively. The CLI, on the other hand, allows automation and scripting for large-scale batch processing. While the GUI aims at assisting individual scientists in their work, the CLI allows the construction of automated workflows in scaled projects. MetaXtract is designed primarily as a Windows-based application, but has been tested and confirmed to work on Linux systems, enabling cross-platform compatibility.

We manually validated that the metadata extracted by MetaXtract matched the information within RAW files by comparing our export with the information accessible in Freestyle version 1.7 (Thermo Fisher). **Figure 1** shows the interface of MetaXtract, which provides an interactive environment for exploring mass spectrometry data. The interface displays key scan parameters such as total ion current, retention time, base peak intensity, and injection time, allowing the user to select all or specific output values for each scan.

**Figure 1.**
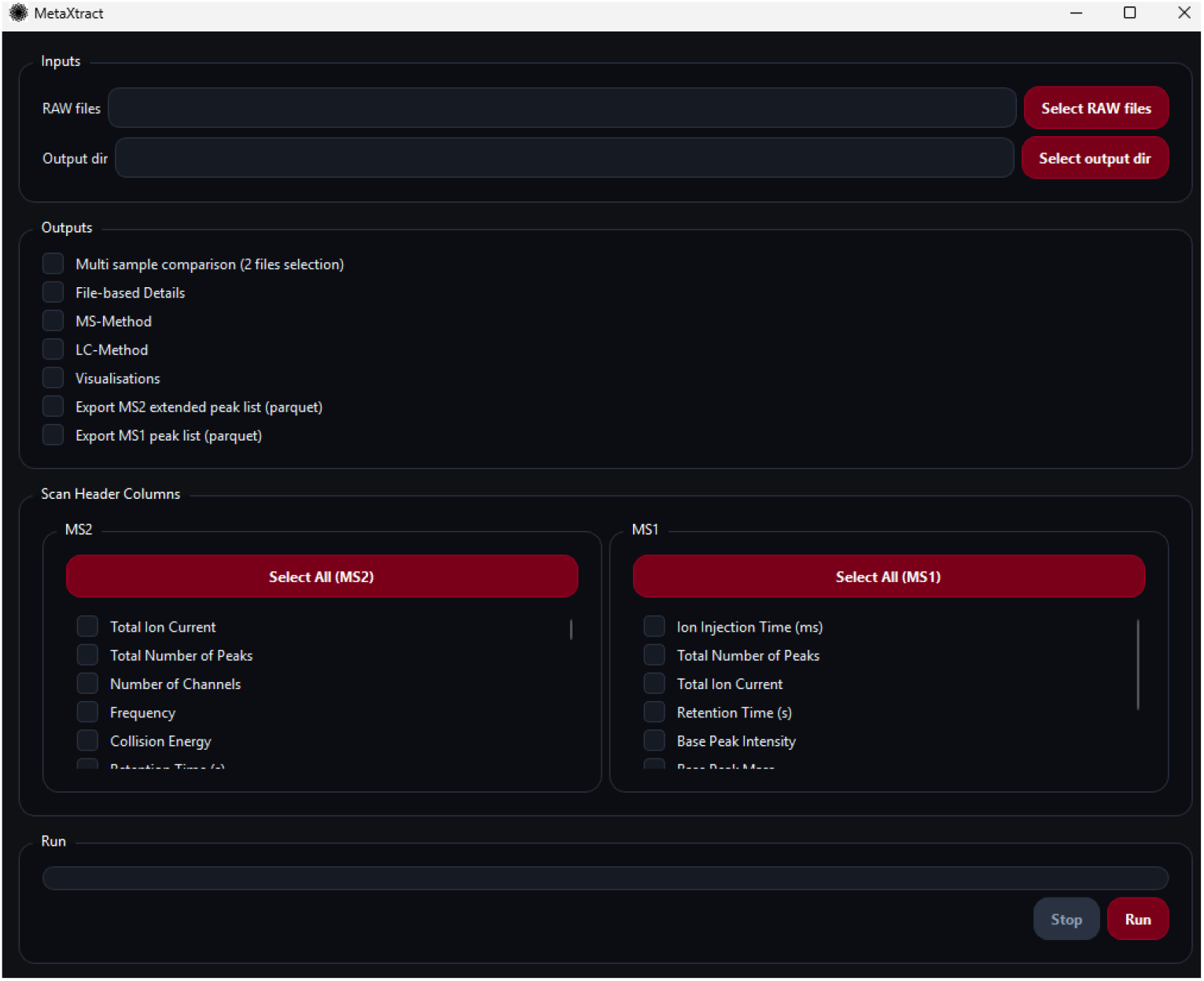
Graphical User Interface of MetaXtract. The interface enables users to select specific mass spectrometry parameters for extraction and visualization. Parameters are grouped into MS1 and MS2 scan information.

### MetaXtract in the context of FAIR data principles

MetaXtract supports several aspects of the FAIR data principles by transforming vendor-specific RAW content into structured, machine-readable output formats such as CSV/TSV or Parquet (for MS1/MS2-extended peaklist). With respect to *findability*, extracted metadata such as instrument settings, scan types, retention time ranges, and sample-related identifiers can be indexed and searched more efficiently than information embedded in proprietary binary RAW files. In terms of *accessibility*, conversion into open, non-proprietary formats reduces dependence on vendor software or operating system-specific libraries, thereby improving access across Linux, macOS, Windows, and cloud-based analysis environments. For *interoperability*, structured tabular outputs, particularly Parquet, facilitate integration with established data science ecosystems such as Python, R, and SQL-based infrastructures, enabling straightforward combination with other omics-derived datasets. Finally, MetaXtract enhances *reusability* by generating reproducible, structured exports together with extraction settings, allowing downstream users to trace how features were defined and processed. In this way, MetaXtract not only simplifies metadata inspection, but also contributes to more transparent and reusable mass spectrometry data handling.

### Workflow integration of MetaXtract to annotate PRIDE submissions

We provide a Snakemake-based workflow in the MetaXtract GitHub repository, fully open source and easily adaptable. It automatically downloads RAW files from PRIDE (by year and file count) and runs our end-to-end annotation. We used this to download the last 20 RAW files submitted to PRIDE in January 2025, spanning separate submissions from three different laboratories (PXD060351, PXD060271 and PXD060112), none being our own. The workflow extracted the full available set of metadata including instrument-, run-, and scan-level metadata (MS1/MS2 headers and method details) into structured TSV/CSV for straightforward reporting. MetaXtract averaged ∼40 s per file on a Windows 11 laptop with 32 GB RAM and NVIDIA GeForce RTX™ 4070 with Intel® Core™ i9 i9-13900HX processor. While we here generate output files, the modular design of MetaXtract allows routing the outputs to downstream tools or LIMS with minimal changes to the workflow.

### Application of MetaXtract for Systematic MS Performance Evaluation

We first demonstrated a possible use case of MetaXtract to monitor mass spectrometer performance independent of database search, enabling near real-time monitoring of data acquisition. We generated comprehensive visualizations of MS1 and MS2 metadata from four datasets of 50 ng HeLa Protein Digest Standard, acquired for quality control purposes. In the two reference runs, 5,055 and 5,843 peptides were identified, whereas only 31 and 5 peptides were identified in the two datasets acquired on a different day, indicating instrument underperformance. The “file-based details” showed a similar total number of MS1 and MS2 scans across both the reference runs (Ref 1, Ref 2) and the underperforming runs (UP 1, UP 2), though the UP runs had a slightly lower proportion of MS2 scans (56% and 59% versus 60% and 62% in the reference runs) (**Figure S3**). Comparable total ion chromatograms (TICs) of MS1 signals suggest that chromatographic separation was not significantly impaired in the UP runs. TIC values for all MS1 scans showed on average 8.4% lower median MS1 signal intensity in the UP runs (**Figure 2a**). Because MS1 behavior remained largely intact while MS2 intensities collapsed, the metadata implicated MS2-specific processes (precursor selection/isolation, fragmentation, or fragment-ion transfer) rather than global LC failure or ionization loss. While this reduction may have contributed to fewer MS2 triggers, it alone is unlikely to account for the dramatic loss in peptide identifications. While protein identifications confirmed the failure post hoc, the MS2 metadata (particularly the strong reduction in MS2 TIC with preserved MS1 TIC) directly pointed to an isolation/fragment transmission problem rather than chromatography. The 9.2-fold reduction in signal intensity in the UP runs (**Figure 2b**) is very likely the primary reason for the loss of peptide identifications. Given the normal performance of MS1 scans, the issue is most likely due to a malfunction during precursor ion isolation or fragment ion transmission. Such near real-time, identification-free quality control (QC) is particularly valuable because it enables early detection of instrument malfunctions or unexpected sample behavior. This is especially critical in high-throughput studies and large-cohort analyses, including single-cell proteomics experiments. By directly leveraging acquisition metadata, our tool enables automated and scalable monitoring of instrument performance and is well suited for near real-time deployment through command-line integration into large-scale workflows. As one application example, MetaXtract can be configured in a pipeline to automatically flag runs where the MS2 TIC drops below a certain percentage of the MS1 TIC, allowing the pipeline to terminate before wasting hours or even days of acquisitions and database searching on underperforming runs.

**Figure 2.**
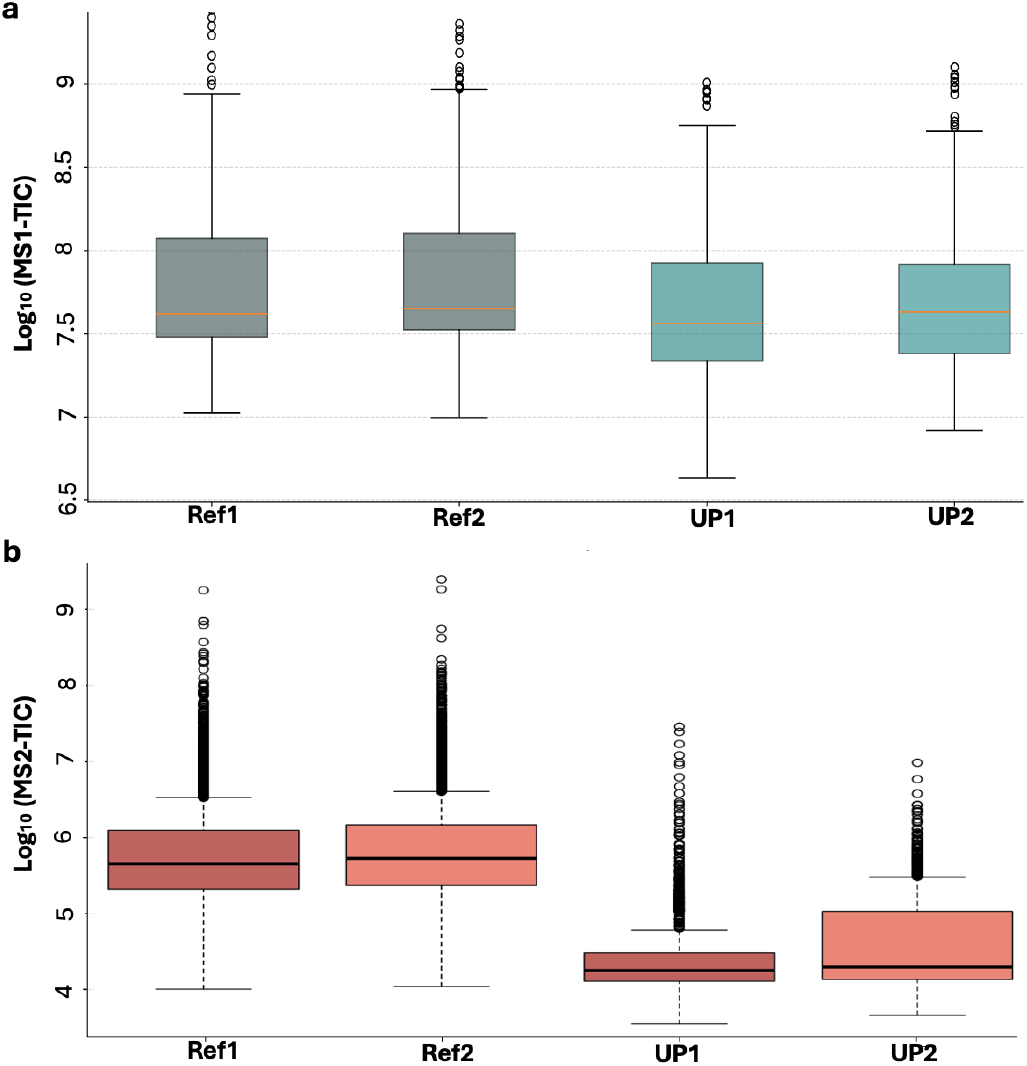
MS1/MS2 total ion current (TIC) across reference and underperforming runs. **(a)** Boxplots of per-scan MS1 TIC reveal lower overall values in the underperforming datasets (UP 1, UP 2) compared with the references (Ref 1, Ref 2). **(b)** Boxplots of per-scan MS2 TIC show a pronounced decrease in UP 2 relative to the references and UP 1, indicating substantial MS2 signal loss and suggesting compromised data quality. *(TIC, total ion current; Ref, reference; UP, underperforming*.)

Beyond metadata, MetaXtract exposes additional run-level parameters useful for routine MS performance tracking and acquisition tuning, illustrated in Supplementary **Figures S4**-**S7**. These metadata can be used in conjunction with downstream identification data, but are also informative on their own. For example, precursor charge state distributions can help monitor digestion efficiency, while plotting MS2 injection times can guide the optimization of maximum ion injection time settings.

## Conclusion

MetaXtract is a lightweight, easy-to-use tool tool for the extraction and visualization of mass spectrometry metadata from Thermo Fisher RAW files. With support for both graphical and command-line interfaces, it offers flexibility for researchers performing manual quality control as well as those integrating it into automated workflows. The command-line functionality allows for simple integration into workflow management systems such as SnakeMake or Nextflow, making it adaptable for high-throughput proteomics pipelines. By combining structured metadata extraction with interactive visualizations of MS1 and MS2 scans and potentially contributing this analysis alongside search results, MetaXtract provides intuitive data exploration, helping researchers optimize acquisition methods and troubleshoot experimental problems. In addition, MetaXtract supports search-free, near real-time QC by enabling immediate evaluation of acquisition-side signals and by linking deviations to plausible instrument-level causes, reducing reliance on delayed identification-based metrics for first-line monitoring. The ability to process multiple RAW files in a single session makes MetaXtract a scalable solution for both small-scale method development and large-scale computational proteomics applications and reinforcing data quality across the field. In future versions, we plan to expand support to additional vendor file formats and integrate MetaXtract as a dedicated node within Proteome Discoverer, further enhancing its utility in diverse proteomics workflows. Beyond its immediate utility for metadata extraction and quality control, MetaXtract also promotes FAIR-oriented data handling by converting proprietary RAW-derived information into structured, searchable, and reusable formats that are accessible across platforms and readily integrated into downstream computational workflows.

## Materials and Methods

MetaXtract is a Python-based mass spectrometry data processing tool (17) designed to support both graphical (built with *PySide6* (18)) and command-line usage. It enables reading, parsing, and visualizing Thermo RAW files by leveraging the Thermo Fisher RawFileReader library (14), which provides direct access to MS1/MS2 scan data, sample metadata, and instrument parameters. The tool adopts a modular architecture, separating data extraction, processing, and visualization into distinct components for improved maintainability and easier integration of future features. The core pipeline includes: (1) RAW file parsing via the *RawFileReader* C# library, (2) data transformation into structured formats (TSV, CSV, PDF), (3) visualization of mass spectrometry plots using Plotly. (4) and export of MS1 and MS2 scan arrays containing all possible information of each scan (Supplementary Algorithms 1-2). The demonstration dataset was acquired using Pierce™ HeLa Protein Digest Standard, 50 ng of Hela digested was used as a quality control (QC) sample was injected for each LC-MS/MS analysis. The LC-MS/MS analysis was performed using a Q Exactive HF mass spectrometer (Thermo Fisher Scientific) connected to an Ultimate 3000 RSLCnano system (Dionex, Thermo Fisher Scientific). Peptides were resuspended in a solution containing 1.6% v/v acetonitrile and 0.1% v/v formic acid, then injected onto a 50-cm EASY-Spray C18 LC column (Thermo Scientific) operating at 50°C column temperature. The mobile phase consisted of water with 0.1% v/v formic acid (mobile phase A) and 80% v/v acetonitrile with 0.1% v/v formic acid (mobile phase B), with peptides loaded and separated at a flow rate of 0.25 μl/min. A gradient was applied, starting with 2% to 9% B over 10 minutes, increasing to 31% B over 30 minutes, further to 44% B over 5 minutes, and finally to 95% B over 2 minutes. Eluted peptides were ionized by an EASY-Spray source (Thermo Scientific) and introduced directly into the mass spectrometer, where MS data was acquired in data-dependent mode. The full scan mass spectrum was recorded with a resolution of 60,000. In each acquisition cycle, the ten most intense ions with a charge state from 2+ to 6+ were isolated and fragmented using higher-energy collisional dissociation (HCD) with a normalized collision energy of 27%. The fragmentation spectra were recorded with a resolution of 15,000. Dynamic exclusion was enabled with a single repetition count and a duration of exclusion of 30 seconds. The data used in this study was acquired on two separate days (19/12/2024 and 20/02/2025, two replicates per date). Protein identification was carried out using MaxQuant (version 2.6.2.0), searching against the human protein sequence database downloaded from UniProt, with the default search parameters.

## Supporting information

PRIDE summary

## Abbreviations

CSV: Comma-Separated Values
TSV: Tab-Separated Values
MS: Mass Spectrometry
QC: Quality Control
LC: Liquid Chromatography
FAIR: Findable, Accessible, Interoperable, and Reusable
LIMS: Laboratory Information Management System

## Acknowledgements

This work was supported by the Wellcome Trust (227434), the Deutsche Forschungsgemeinschaft (DFG, German Research Foundation) under Germany’s Excellence Strategy - EXC 2008 - 390540038 - UniSysCat and the European Research Council (ERC) under the European Union’s Horizon Europe research and innovation programme (grant agreement No. 101119142 - TransFORM).

## Supplementary Materials

### Figures

**Figure S1.**
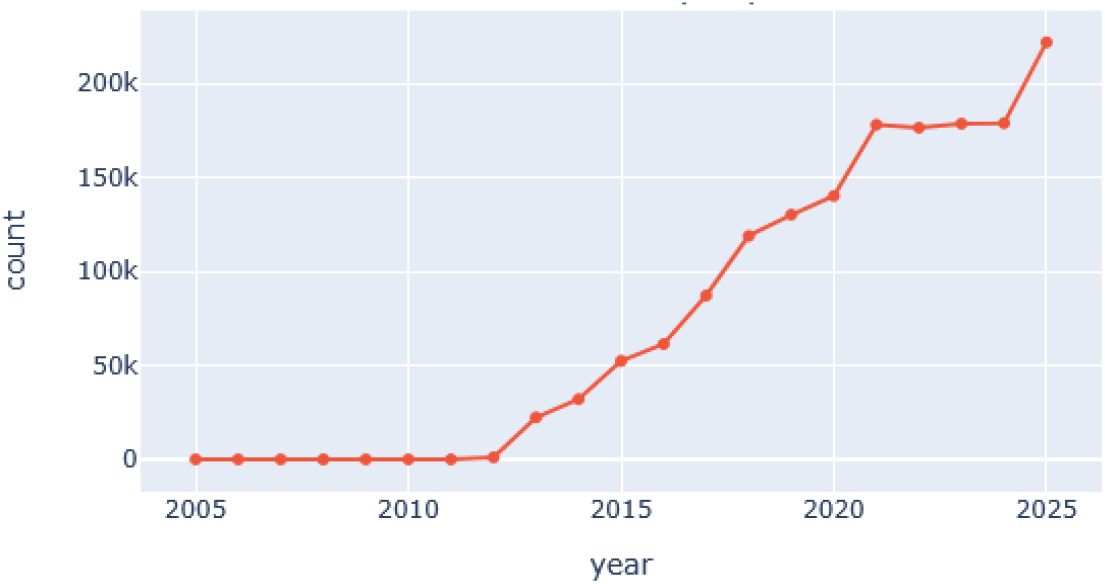
Annual growth in RAW file submissions (2005-2025). The number of submitted RAW files remained negligible until approximately 2012, followed by a sustained and steep increase from 2013 onward. Submissions rose sharply between 2015 and 2019, exceeded 150,000 annually by 2021, and reached approximately 220,000 by 2025. The trend reflects rapid expansion in mass RAW files generation and submission over the past decade.

**Figure S2.**
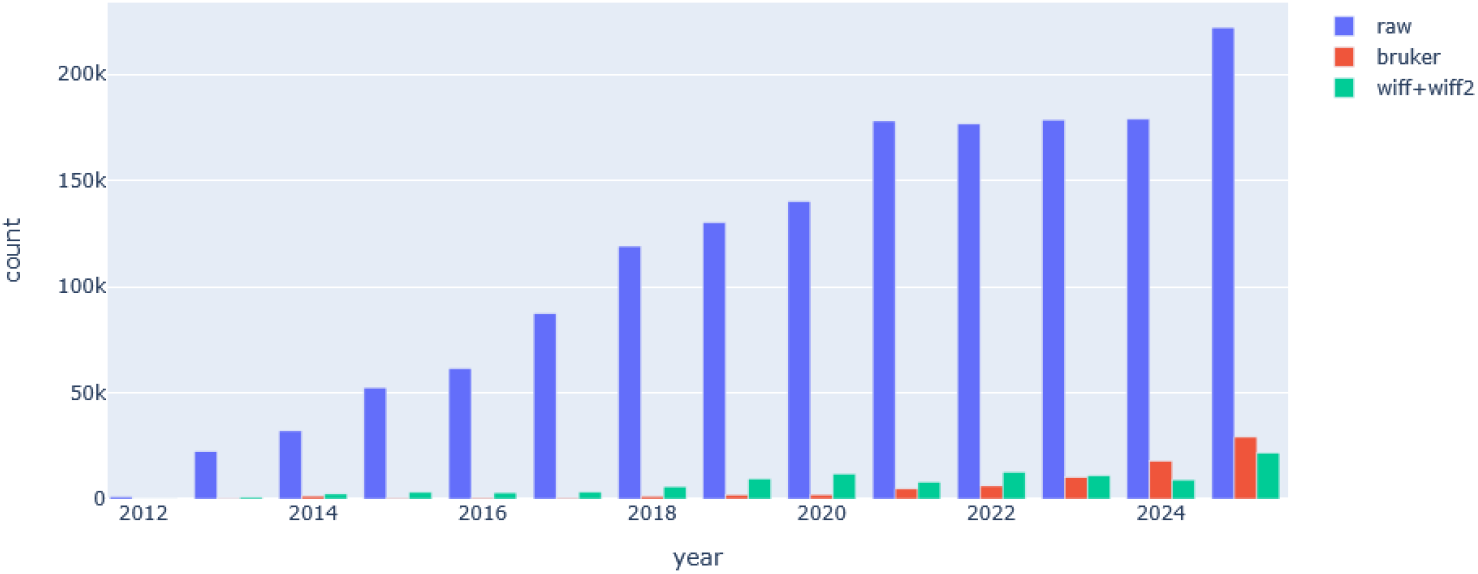
Annual submissions by MS file format (2012–2025). Thermo RAW files dominate public submissions across all years, exhibiting rapid growth from 2012 onward and exceeding 200,000 annual entries by 2025. Bruker formats show a delayed but marked increase after 2020, with substantial growth through 2024–2025. WIFF/WIFF2 submissions rise steadily over time but remain lower in absolute numbers compared to Thermo and Bruker formats.

**Figure S3.**
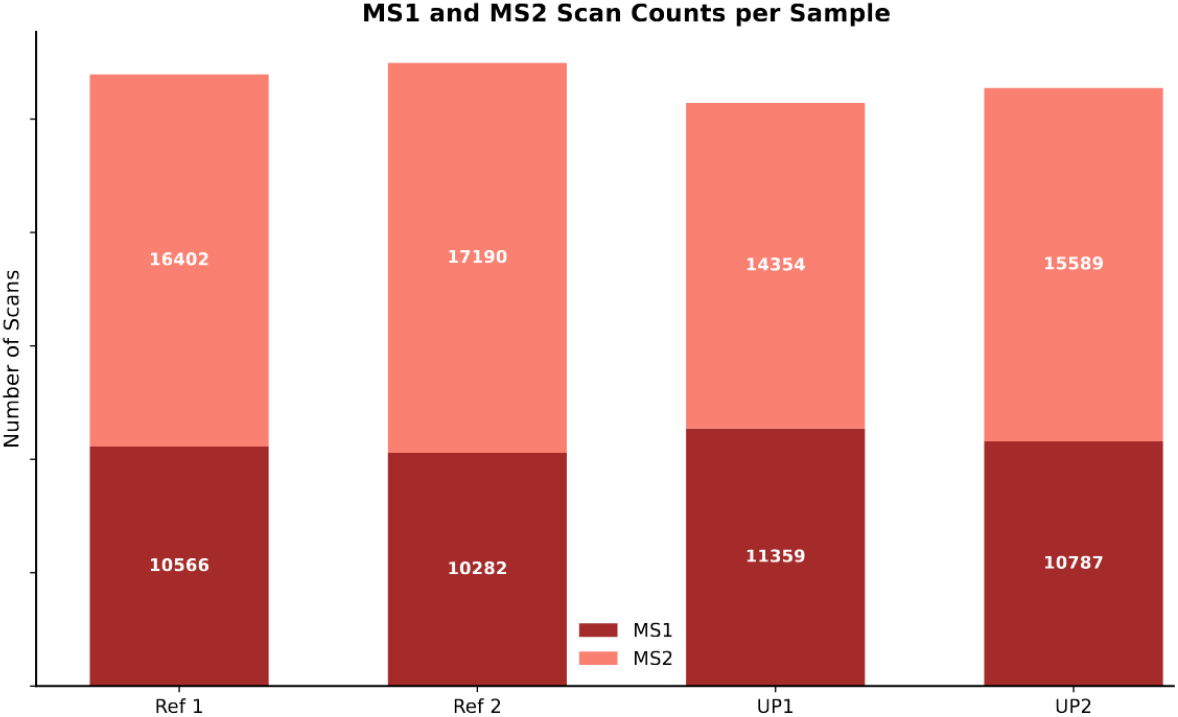
MS1 and MS2 scan counts across samples. The bar plot shows the number of MS1 and MS2 scans for four samples. Minor variations in scan counts reflect run-specific acquisition behavior, highlighting the importance of comparative scan-level QC.

**Figure S4.**
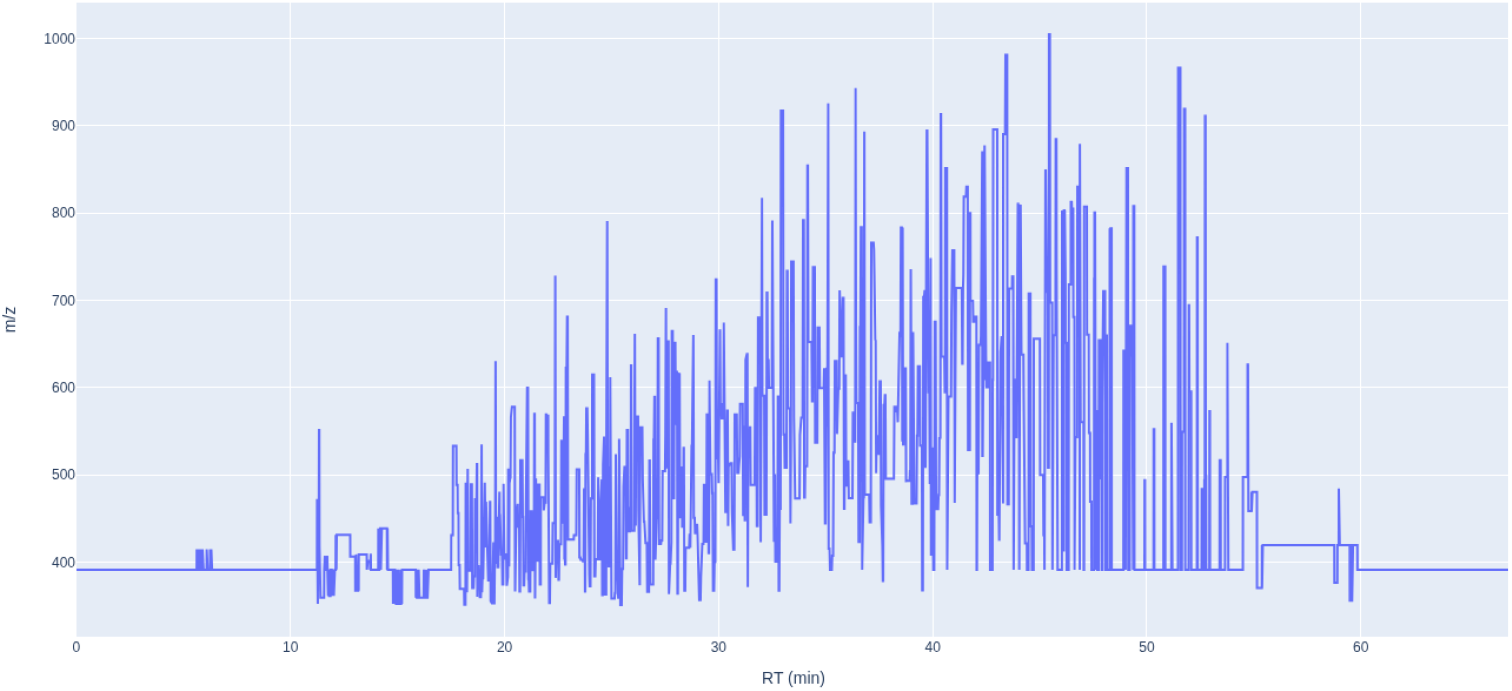
Base-peak mass vs. retention time for the first reference run, showing the distribution of detected base peaks across the gradient and a mild shift toward higher base-peak mass at later retention times.

**Figure S5.**
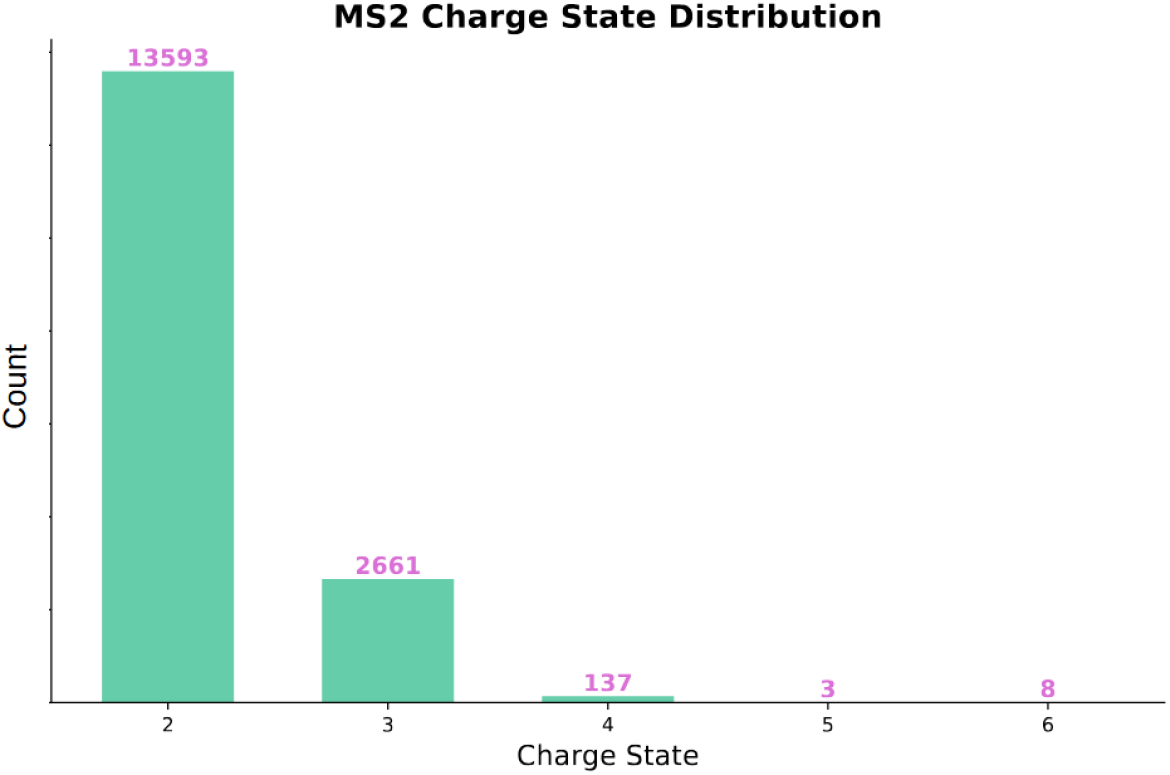
Charge-state distribution for the first reference run.

**Figure S6.**
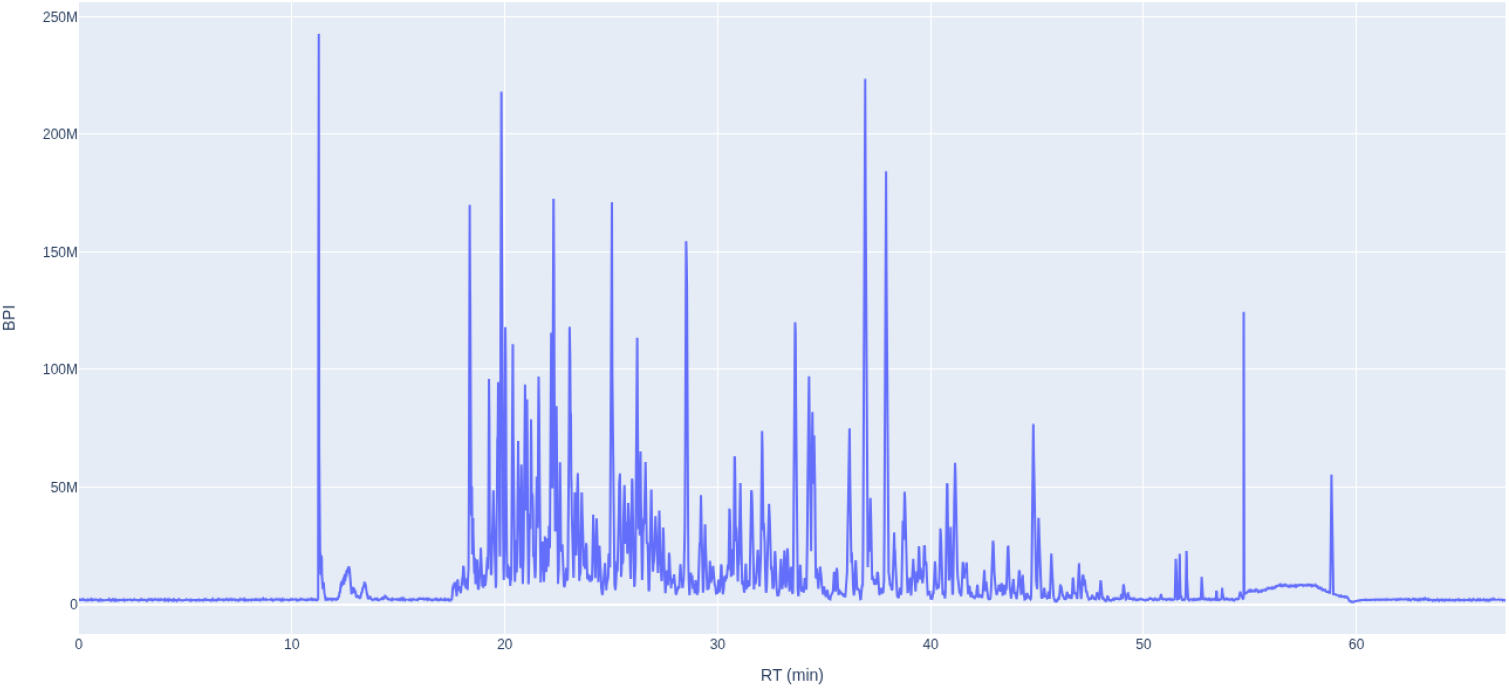
Base-peak intensity vs. retention time for the first reference run, showing the main elution window (∼11.20-58 min) with dense apexes around 18-45 min and a low baseline outside; sharp spikes mark isolated high-abundance features.

**Figure S7.**
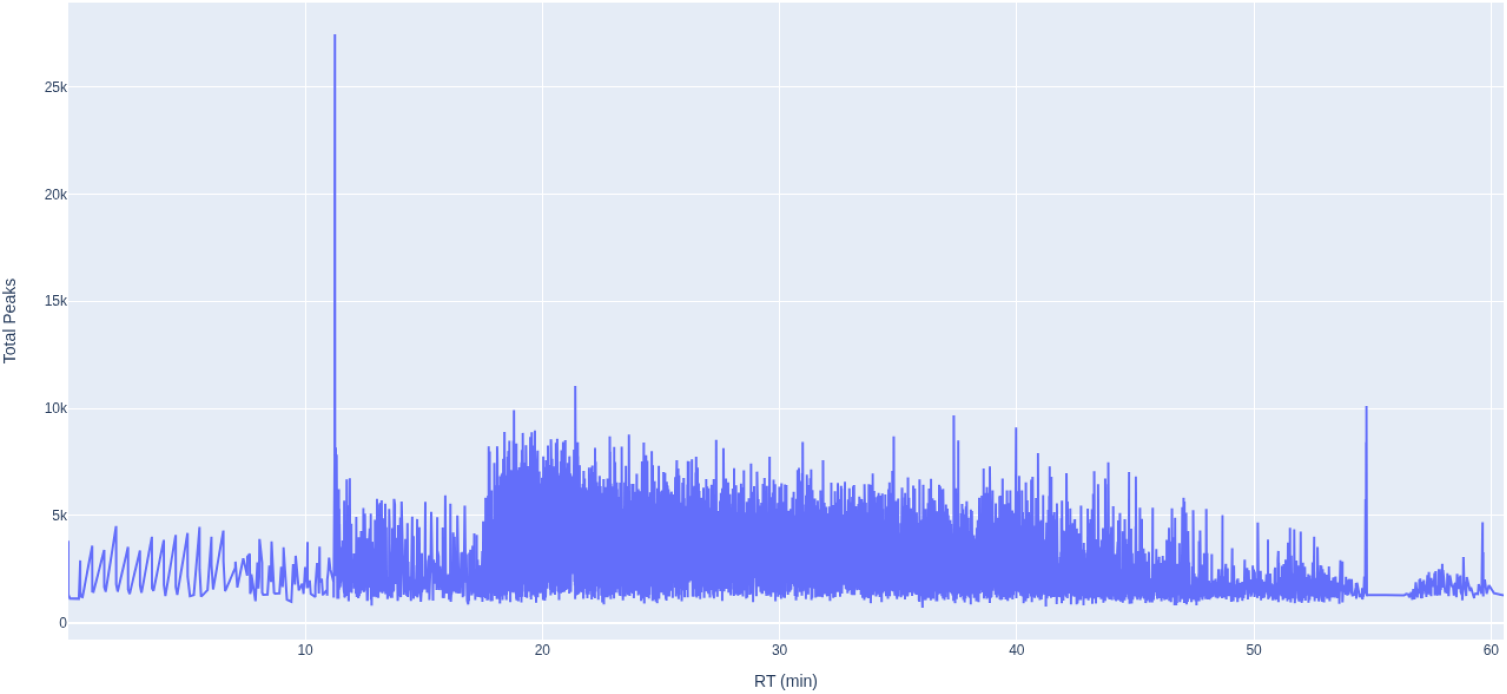
Total number of MS2-peaks vs. retention time for the first reference run, spike at ∼11.20 min, broad high-density elution from ∼12-53 min with a gradual decline, and low activity outside the window.

### Algorithms

This supplement provides high-level pseudocode for the peak list extraction and export routines

#### Algorithm 1

MS2 peak list extraction (per scan)

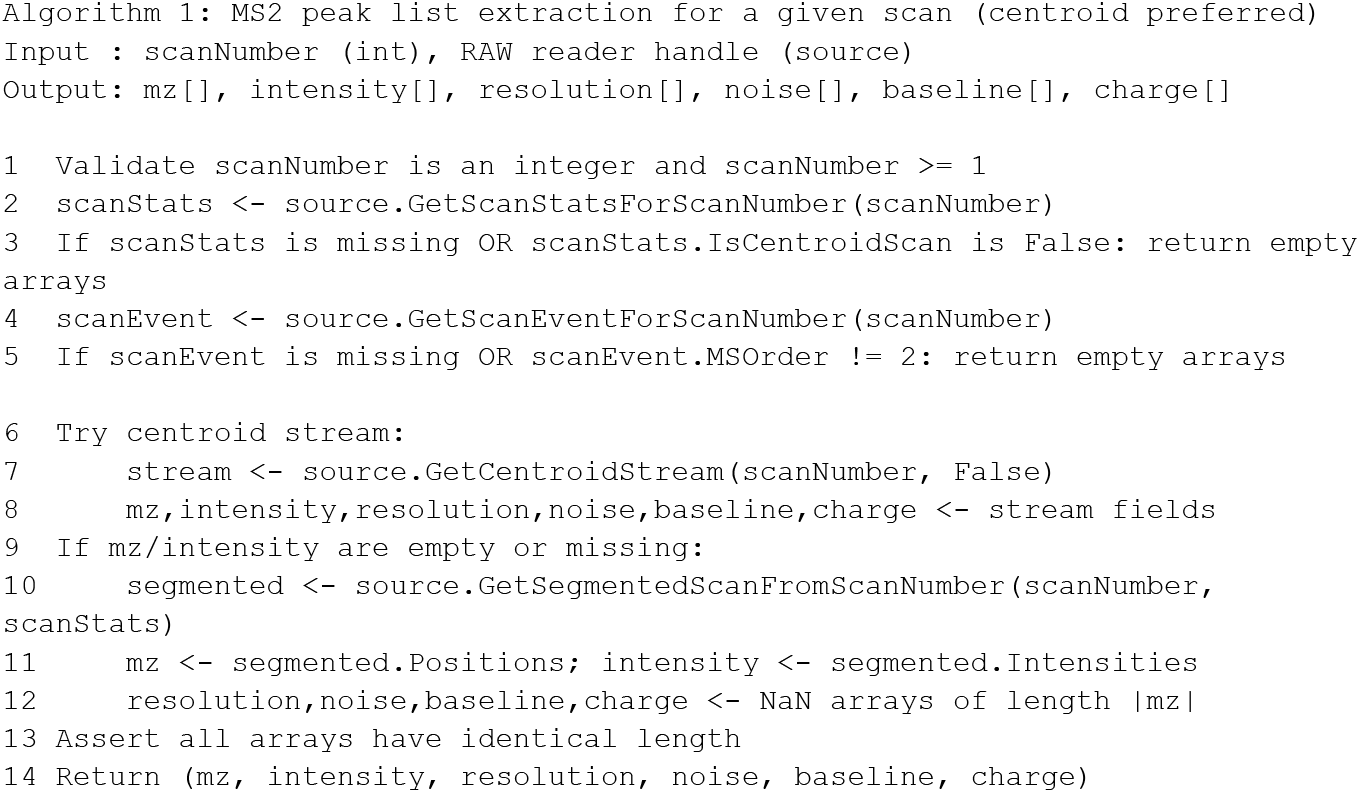

#### Algorithm 2

MS1 peak list export (chunked Parquet)

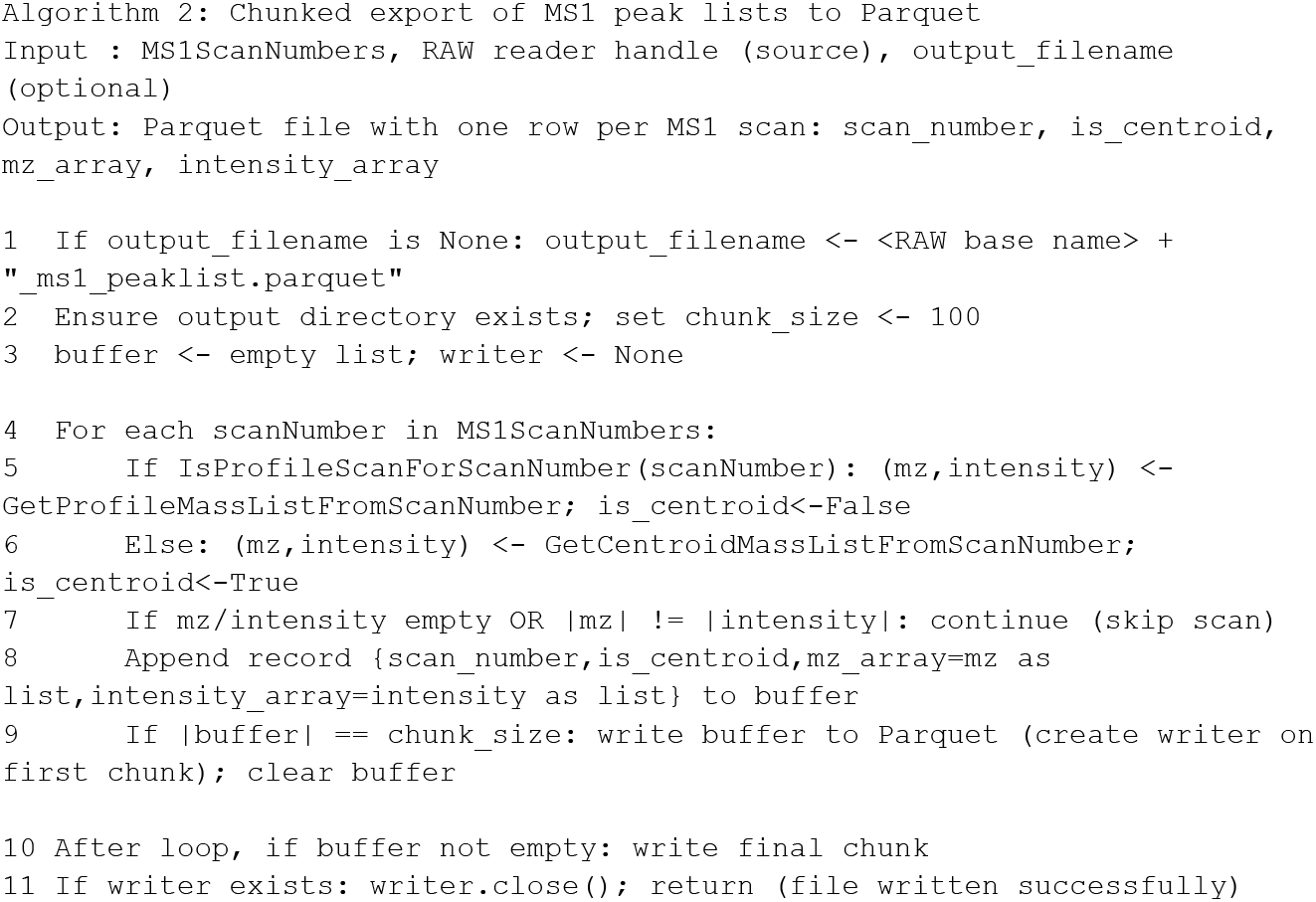

## Notes

### Competing Interest Statement

The authors have declared no competing interest.

### Summary of Updates

Tool changes and code optimisations. Some aspects are changed in the paper describing the method applied.

