## Supplementary material for "MetaXtract: Extracting Metadata from Raw Files for FAIR Data Practices and Workflow Optimisation": PRIDE summary: summary_pride.html

PRIDE MS filetypes by year (2000–2025)


### PRIDE MS filetypes by year (2000–2025)

#### Summary

**MS data (including peak list):**

- **raw** : Thermo RAW; vendor raw data
- **d** : Bruker “.d” dataset directory; vendor raw data container
- **baf** : Bruker BAF; vendor raw data
- **tdf** : Bruker timsTOF raw data files
- **tdf\_bin** : Bruker timsTOF raw data files
- **wiff** : SCIEX vendor raw data
- **wiff2** : SCIEX vendor raw data
- **mzml** : open raw spectra format
- **mzxml** : open raw spectra format
- **mgf** : peak list / MS2 spectra export

##### MS data table
